# Cholinergic interneurons in the dorsal striatum play an important role in the acquisition of duration memory

**DOI:** 10.1101/2023.08.18.553866

**Authors:** Masahiko Nishioka, Toshimichi Hata

## Abstract

Although the formation of duration memory is important to optimize the timing of behavior based on previous experiences, the neural mechanism of formation remains unclear. A previous study suggested that muscarinic acetylcholine 1 receptors (M1Rs) in the dorsal striatum of rats are involved in the consolidation of duration memory in interval timing. Therefore, cholinergic interneurons (ChIs) may also be involved in the formation of duration memory in interval timing because ChIs activate M1Rs in the dorsal striatum. In the Exp. 1A, rats underwent a peak interval (PI)-20 s training. During the training, two different trials were randomly presented. One was a food trial in which the initial lever press was reinforced 20 s after the start of the trial (i.e., fixed interval (FI), 20 s). The other was an empty trial in which the reinforcement did not occur for 80 s. After sufficient training, the rats responded at approximately 20 s in the empty trials. They were then subjected to a PI-40 s training (i.e., FI, 40 s) after ChI lesions were present in the dorsal striatum. In this training, the sham-lesioned group responded frequently at approximately 40 s, whereas the ChI-lesioned group responded at approximately 20 s. As the PI-40 s additional learning progressed in the Exp. 1B, the ChI-lesioned group frequently responded at approximately 40 s, similar to that of the sham-lesioned group. In the following PI-20 s re-shift training, the ChI-lesioned group responded similar to the sham-lesioned group. In Exp. 2 of another cohort, the results of a PI-20 s training after the occurrence of ChI lesions were similar to that before the presence of the lesion. Together, these results suggest that ChI lesions delayed only the acquisition of new duration memory, but had no effect on the adjustment of behavior associated with changing the reinforcement schedule of the PI-training and interval timing itself.

## Introduction

Interval timing is the ability to perceive durations ranging from seconds to minutes and modulate behavior based on it; it is required for organisms to forage and is responsible for their behavior (Buhushi & Meck, 2005). To perform interval timing, the formation of duration memory is necessary. Data on the peak-interval (PI) procedure (Roberts, 1981) suggested that animals can form duration memory. In this procedure, food trials and empty trials were presented in a random order. Both trials started with an onset of signal stimuli and ended with an offset of them. In the food trials, the initial lever-press was reinforced after a predetermined time *t* (a required time) from the beginning of the trial (*i.e*., discrete fixed-interval (FI) schedule). In the empty trials, no responses were reinforced even after the required time. After sufficient training, rats finally began pressing the lever just before the time *t* and stopped pressing the lever after the time *t* elapsed in empty trials. This behavioral pattern showed that rats can form duration memory and optimize the timing of their behavior using this memory. However, the neural mechanism of the formation of duration memory remains unclear. Thus, it is important to clarify this mechanism to better understand the mechanism of interval timing.

Muscarinic acetylcholine 1 receptors (M1Rs) in the dorsal striatum (DS) are essential for the consolidation of duration memory (Nishioka et al., 2022). In this study, rats underwent a PI-20 s training (*i.e*., food trials had a FI of 20 s). After sufficient training, the response distribution as a function of the elapsed time in an empty trial was depicted as curves with peaks around 20 s. To make the rats acquire new duration memory, they were subjected to a PI-40 s training after injecting an M1R antagonist, pirenzepine, into the DS. In this training, the peaks of curves in the control and M1R-inhibited groups similarly shifted toward 40 s. A test session with no drugs was conducted a day after the shift session, and only empty trials were presented. In the test session, the peak of the control group was at approximately 40 s, whereas that of the M1R-inhibited group was at approximately 20 s, similar to those of the training sessions before changing the required time. These results suggest that injection of the M1R antagonist into the DS impaired the consolidation of duration memory, which lasted until the next day. This interpretation is consistent with an operational definition of the impairment of consolidation, that is, an impairment of the retention of an acquired memory under the influence of a drug until the next day (Johansen et al., 2011). Therefore, it is suggested that M1Rs in the DS are essential for the consolidation of duration memory.

Cholinergic interneurons (ChIs) in the DS may also be involved in the formation of duration memory through input to M1Rs in medium spiny neurons (MSNs). Inputs from ChIs are a major source of acetylcholine in the DS and are involved in increasing the excitation of MSNs (Narushima et al., 2007; Goldberg & Reynolds, 2011). Physiological studies on temporal reward prediction in monkeys showed that tonically active ChIs in the DS “paused” following the presentation of an antecedent stimulus of reward and the reward itself (Apicella et al., 1991; Apicella et al., 1997; Apicella et al 2009; Martel & Apicella, 2021; Martel & Apicella, 2022). For example, the pause in an early phase of training was stronger than that in the late phase (Sardo et al., 2000; Ravel et al., 2001), showing that the pause especially occurs in a novel temporal association learning. The presentation of a reward at a predictable timing after sufficient amount of conditioning resulted in a weak pause, whereas the presentation of a reward at an unpredictable timing resulted in a strong pause (Revel et al., 2001). These characteristics may play a role in the formation of new duration memory. Therefore, it is suggested that ChIs in the DS may also be involved in the formation of duration memory. However, this possibility has not been operationally investigated.

The present study has investigated the effect of ChI lesions in the DS on the formation of duration memory using the same behavioral paradigm as the previous work (Nishioka et al., 2022). Rats received PI-20 s trainings, followed by PI-40 s trainings under ChI lesions. As a result, peaks of the response curve of the aCSF group shifted from 20 s to 40 s, whereas that of the ChIs group remained at approximately 20 s and was similar to those of trainings before the shift training. These results suggest that ChIs in the DS play an important role in the acquisition of duration memory.

## Exp. 1A

The purpose of this experiment was to clarify the effect of ChI lesions in the DS on the acquisition of duration memory. All experimental procedures were approved by the Animal Research Committee of Doshisha University (A22085).

## Materials & Methods

### Subject

Twenty-two naïve male albino Wistar rats aged 3 months old were included in this experiment. They were individually housed in cages. Before the training, we limited their body weight to 90% of their free-feeding weight to increase the drive for food. Considering their growth, we then increased their weight limits by 5 g every week during training. Water was provided *ad libitum* in the cages. The breeding room was lighted from 8:00 AM to 8:00 PM, and the experiment was conducted during the light phase.

### Apparatus

Eight identical operant chambers were used. Each of them was equipped with a lever (40 mm above the floor) to the right of a food cup (15 mm above the floor) at the center of a front wall. The walls and ceiling of the chamber were made of acrylic board and the floor was made of stainless bars (3 mm diameter) separated by 10 mm. These chambers were individually placed in a sound-proof box equipped with a 0.8 W LED and a buzzer (M2BJ-B24, OMRON, Kyoto, Japan) on the ceiling. An application to control the experimental devices and record data was developed using XOJO® (XOJO Inc. Austin, TX, USA) on a MacBook Air (Apple, Cupertino, CA, USA). A programmable controller (SYSMAC CPM1A40CDR-AV1, OMRON, Kyoto, Japan), a keyboard encoder (Bird Electron, Kanagawa, Japan), and a USB I/O controller (RBIO-2 U, Kyoritsu Electronic, Osaka, Japan) were interfaced with the MacBook. The system could record the lever-press time with 0.1 s temporal resolution.

### Procedure

#### Surgery

After 5 min of handling the subjects per day for five days, the subjects were anesthetized with 2–3% isoflurane by inhalation (2–3 L/min flow rate, MK-A110, Muromachi, Tokyo, Japan) and fixed on a stereotaxic frame (David Kopf Instruments, Tujunga, CA, USA). The tip of a bilateral guide cannula (C232G-5.0/SPC, Plastics One, Roanoke, VA, USA) were placed in the DS (from bregma, AP: +0.5 mm, ML: ±2.5 mm, DV: -3.8 mm from the skull surface). Note that artificial cerebrospinal fluid (aCSF) or immunotoxin was not injected here. The cannula was fixed using dental cement and three small screws on the skull. The cannula was plugged with a dummy cannula (C232DC-5.0/SPC, Plastics One) and screwed with a dust cap (363DC, Plastics One). Antibiotics (Mycillinsol, Meiji, Tokyo, Japan) were applied to the injured area.

#### Shaping

A week after the surgery, the subjects were placed into the chambers for 10 min for habituation. Next, they received lever-press trainings by a continuous-reinforcement schedule under LED illumination. In this training, a lever press was reinforced by a food pellet (F0021-J, Flemington, NJ, USA). The acquisition criterion was the pressing the lever 60 times within 20 min in three successive sessions.

#### Training (PI-20 s, sessions 1–30)

The subjects had one PI-20 s training session per day. Food and empty trials were presented in this training. Both trials started with an onset of the light and buzzer and terminated with an offset of both. In the food trials, the initial lever press 20 s (i.e., required time) after starting the trial produced a pellet and marked the end of the trial. In the empty trials, none of the lever-presses resulted in a reinforcement, and the trial was terminated 80 s from the beginning of the trial. Both trials were presented in a pseudo-random sequence in a way that prevented the presentation of four consecutive empty trials. The trials were separated by inter-trial intervals (ITIs) of 40 ± 10 s. When the lever was pressed within 5 s before the scheduled termination time, the ITI was extended for 10 s. The ratio of food to empty trials was 42:18 in sessions 1–20 and 15:15 in sessions 21–30 (Fig. 2A).

#### Immunotoxin injection

The subjects received an immunotoxin injection a day after the 30^th^ session (Fig. 2A). The subjects were divided into two groups so that the peak times and CVs through the last three sessions of the training would be even. Under anesthesia, in the same way as the surgery, the dummy cannula was replaced with an injection cannula (C232I-5.0/SPC, Plastics One). The tip of the injection cannula extended 1 mm beyond the tip of the guide cannula. Each group of rats had either aCSF or anti-choline acetyltransferase (ChAT)-saporin (ChAT-SAP, IT-42, Advanced Targeting Systems, California, USA) infused into the bilateral DS using a gas-tight syringe (100 μL, Hamilton, Reno, NV, USA) and a micro-syringe pump (0.5 μL/min of flow rate, ESP-32, Eicom, Kyoto, Japan) for 1 min per side. The concentration of the ChAT-SAP was 0.5 μg/μL. The injection cannula was left in place for 1 min after the injection to allow dispersion of the solution. The injection cannula was then replaced with the dummy cannula.

#### Shift training (PI-40 s, session 31)

To make the subjects acquire new duration memory, we subjected them to a PI-40 s training (i.e., required time, 40 s). This training was conducted 72 h after aCSF or ChAT-SAP injection because saporin-mediated cell death occurs within this period. The other parameters used are the same as in the 30^th^ session (Fig. 2A).

#### Test (session 32)

To confirm whether the memory of duration had been consolidated, we tested them a day after the shift training. Note that this session consisted of 15 empty trials only. The other parameters were the same as in the 30^th^ session (Fig. 2A).

### Histological method

After each experiment, the subjects were anesthetized with a lethal dose of sodium pentobarbital (130 mg/kg, i.p.) and perfused with osmotic pressure-regulated phosphate buffer (PB) (sucrose 8.5% in 20 mM PB), followed by 200 mL of 4% paraformaldehyde (PFA) and 0.2% picric acid in 0.1 M PB solution. Next, the extracted brains were postfixed in the same PFA solution for 3 h. The brains were immersed in 15% and 30% sucrose followed by 0.1 M PB until they sunk in each solution at once and twice, respectively. We prepared 40 μm coronal sections using a cryostat (CM1850, Leica, Wetzlar, Germany) for immunohistochemistry. The sections were incubated overnight by polyclonal anti-ChAT goat antibody (AB144P, Merck millipore, MA, USA), followed by a 3-h incubation with polyclonal Donkey anti-Goat Alexa flour 488 antibody (AB 2340428, Jackson immunoresearch, PA, USA). These antibodies were diluted by ×500. Images of the sections were taken at ×4 magnification using a microscope (Eclipse 80i, Nikon, Tokyo, Japan).

### Data processing

The parameters of lever-press behavior from empty trials were used for analysis.

#### Session-by-session analysis

The total number of lever-press responses in 3-s bins (such as 0–3 s and 1–4 s) in each session were counted individually. These bin data were converted into percentages by dividing the number of responses of each bin by the maximum number of responses for all bins in each session. These values were fitted by the gaussian curve (*R(t) = a + b exp{-(t-c)/d}*^*2*^) using the fitnlm function (MATLAB, ver. R. 2020a, MathWorks, MA, USA). *R(t)* shows the estimated response rate at time *t*; *c* was defined as a peak time that shows the subjective length for the required time and duration memory. The initial values of *a, b*, and *d* were set to 0, 100, and 8, respectively. The initial value of *c* was the mean of the middle values of the two bins, the first and last bins having response rates of over 85%. To converge the fitting results of the shift training, we omitted data of the first five trials from the fitting. To estimate the precision of the peak time and the motivation for reward, we calculated the coefficient of variance of the peak time ((*d/*√*2)/peak time*) and the peak rate (*maximum number of responses in all bins/number of trials that were included in session-by-session analysis*). The 30^th^ session was defined as the baseline.

#### Trial-by-trial analysis

The start and stop times in each empty trial were calculated. In the empty trials, the subjects typically started pressing the lever before the required time and stopped pressing it when the required time elapsed. This behavioral pattern was called “low-high-low” pattern. The changepoints from low to high and high to low states were defined as the start and stop times, respectively. These indices were used as a subjective length for the required time and duration memory. These points were calculated using an individual trial analysis (Church et al., 1994) and were estimated by the method of previous studies (Lusk et al., 2020; Yousefzadeh et al., 2021) using Matlab. Briefly, the total lever presses in 1-s bins in each trial were counted indivually. Ten zeros were then added to the beginning and end of the bin data in every trial. The padded bin data were processed using a median filter (window size = 5), followed by a sgolayfilter (sgolayfilt function; window size = 3) and log-normalized. The two changepoints where the residual error was minimum were estimated using the “findchangepts” function. The difference between the stop time and start time in each trial was defined as a width. Trial data with only one or no changepoint were excluded from the analysis.

### Statistical analysis

Two-way mixed ANOVA was used. Multiple comparisons were conducted using Holm’s method. The α value was .050. When the sphericity assumption was violated, the degree of freedom was adjusted using Greenhouse-Geissers’ ε.

## Results

### Histological results

Figure 1 shows representative sections of the DS (A) and the extent of the lesion (B). A rat died before perfusion and was not included in Figure 1B. ChIs were observed near the track of the guide cannula in a section of the aCSF group, but not in the ChAT-SAP group (Fig. 1A). The maximum and minimum lesion areas were limited within the DS (Fig. 1B).

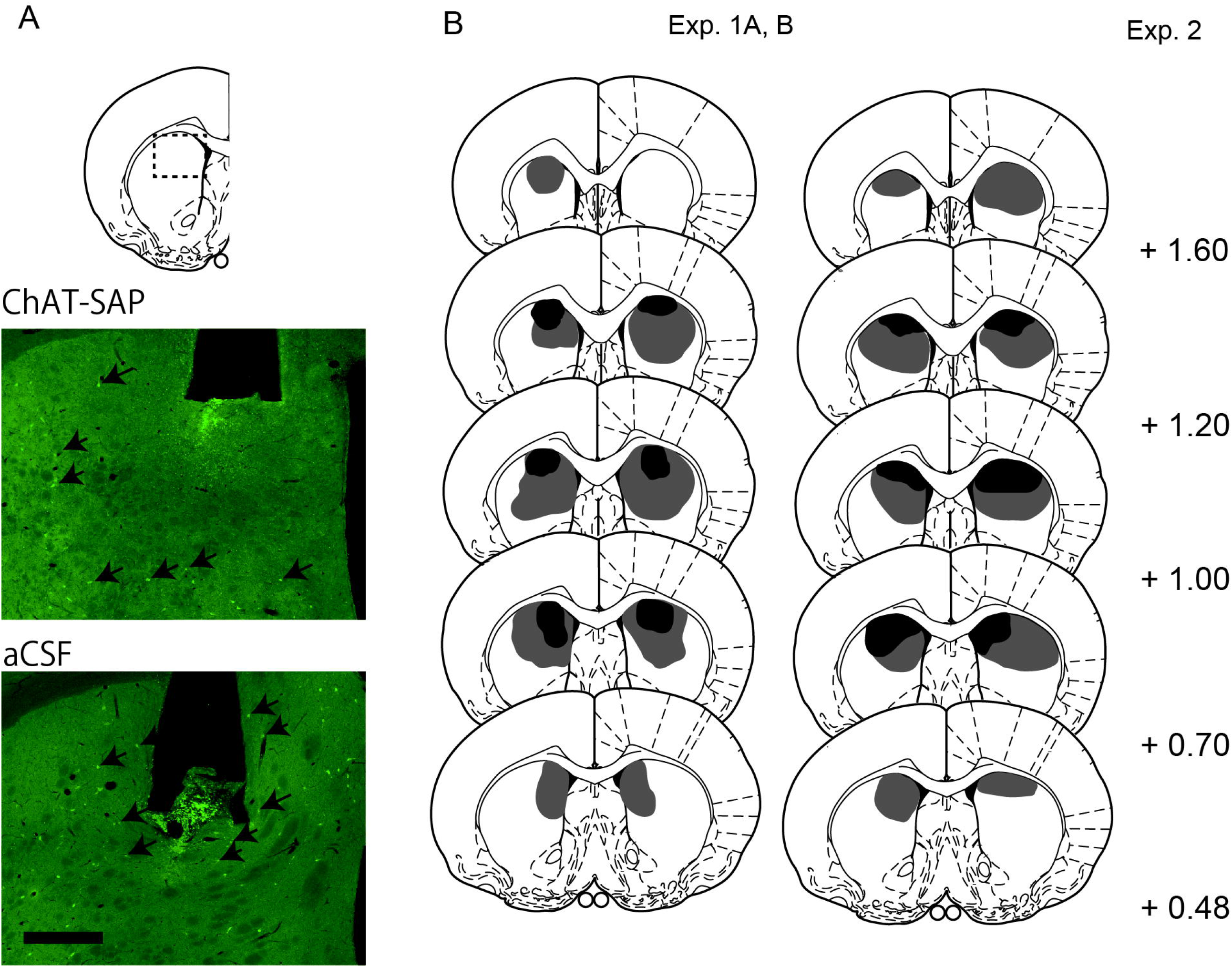

### Behavioral results

Data from four rats were not included in the analysis of Exp. 1A. Two were because of failure to converge for fitting, one was because of negative value of an estimated peak time, and one were due to poor *R*^*2*^s of fitting curves (below -2*SD*). Thus, the data of 10 and eight subjects were included for the ChAT-SAP and aCSF group, respectively. The data of one rat in the ChAT-SAP group that died before perfusion were included in the analysis because the lesion was assumed to be similar to that of the other rats.

### Response distribution

Figure 2B shows mean response-rate distributions from the baseline to the test. At the baseline, the distributions of both groups overlapped with peaks at approximately 20 s (Fig. 2B, left). In the shift training, the peak of the distribution of the aCSF group shifted to approximately 30 s, whereas that of the ChAT-SAP group remained at approximately 20 s (Fig. 2B, middle). In the test, the peak of the distribution of the aCSF group remained at approximately 30 s, whereas that of the ChAT-SAP group remained at approximately 20 s (Fig. 2B, right).

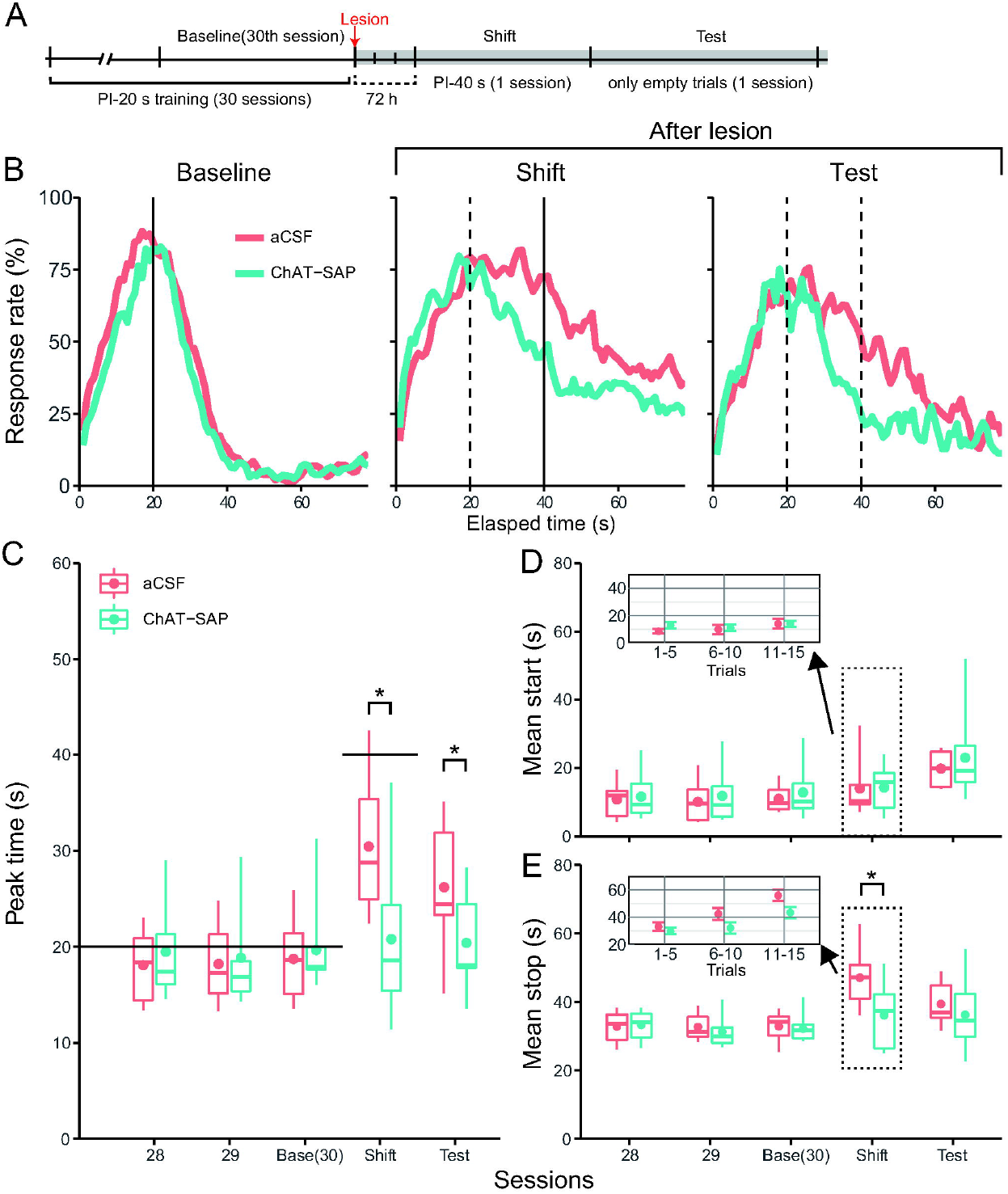

#### Peak time

Figure 2C shows a box and whisker plot of peak times from the 28^th^ session to the test. The peak times of both groups were consistently at approximately 17 s from the 28^th^ session to the baseline (Fig. 2C). In the aCSF group, the peak time of the shift training was higher than that of the baseline, whereas the peak time of the ChAT-SAP group was similar to that of the baseline and lower than that of the aCSF group. In the test, the peak times of both groups remained at values similar to that of the previous session. The mixed two-way ANOVA (group (2) × session (5)) showed that the main effect of the session and the interaction were significant (session: *F*(2.41, 38.54) = 17.964, *p* < .001, η*G*^*2*^ = .229; interaction: *F*(2.41, 38.54) = 10.821, *p* < .001, η*G*^*2*^ = .152), but not the main effect of the group (*F*(1, 16) = 1.227, *p* = .284, η*G*^*2*^ = .053). The simple main effects of the group in the shift training, test (shift: *F*(1, 16) = 7.273, *p* = .016, η*G*^*2*^ = .313; test: *F*(1, 16) = 4.672, *p* = .046, η*G*^*2*^ = .226), and session in the aCSF group (*F*(1.44, 10.11) = 21.116, *p* = .001, η*G*^*2*^ = .502) were significant. Multiple comparisons of sessions in the aCSF group showed that the values in the shift training and test were significantly higher than those in the other sessions (*p*_s_ < .045).

#### Start time

Figure 2D shows a box and whisker plot of mean start times from the 28^th^ session to the test. The values of both groups increased similarly in the test (Fig. 2D). The mixed two-way ANOVA showed that the main effect of the session was significant (*F*(1.93, 30.88) = 16.685, *p* < .001, η*G*^*2*^ = .243), but not the main effect of the group and interaction (group: *F*(1, 16) = 0.309, *p* = .586, η*G*^*2*^ = .013; interaction: *F*(1.93, 30.88) = 0.260, *p* = .765, η*G*^*2*^ = .005). Multiple comparisons of sessions showed that the values in the test were significantly higher than those in the other sessions (*p*_s_ < .017). The inset in 2D shows the mean median of start times (*±SEM*) of every five trials in the shift training. The values of both groups were similar in all trials. The mixed two-way ANOVA (group (2) × trial (3)) showed that the main effect of the trial was significant (*F*(2, 32) = 3.871, *p* = .031, η*G*^*2*^ = .046), but not the main effect of the group and interaction (group: *F*(1, 16) = 0.326, *p* = .576, η*G*^*2*^ = .016; interaction: *F*(2, 32) = 1.264, *p* = .296, η*G*^*2*^ = .015). However, multiple comparisons of the trial showed that none of the pairs were significant.

#### Stop time

Figure 2E shows a box and whisker plot of mean stop times from the 28^th^ session to the test. The values of both groups were similar in the PI-20 s trainings (Fig. 2E). In the shift training, the value of the ChAT-SAP group was lower than that of the aCSF group and similar to that at the baseline. In the test, the values of both groups were similar. The mixed two-way ANOVA showed that the main effect of the session and interaction were significant (session: *F*(2.64, 42.31) = 11.323, *p* < .001, η*G*^*2*^ = .276; interaction: *F*(2.64, 42.31) = 3.391, *p* = .031, η*G*^*2*^ = .103), but not the main effect of the group (*F*(1, 16) = 2.338, *p* = .146, η*G*^*2*^ = .063). There were significant simple main effects of the group in the shift training (*F*(1, 16) = 6.407, *p* = .022, η*G*^*2*^ = .286) and of the session in the aCSF group (*F*(2.10, 14.72) = 17.528, *p* < .001, η*G*^*2*^ = .532). Multiple comparisons of sessions in the aCSF group showed that the values in the shift training were higher than those in the other PI-20 s training sessions, and that the value in the test was higher than those from the 28^th^ session to the baseline (*p*_s_ < .012). The inset in 2E shows the mean and median of start times (*±SEM*) of every five trials in the shift training. The values in the ChAT-SAP group were lower than that of the aCSF group in the 6^th^ and 11^th^–15^th^ trials. The mixed two-way ANOVA showed that the main effects of the group and of the trial were significant (group: *F*(1, 16) = 5.154, *p* = .037, η*G*^*2*^

= .174; trial: *F*(2, 32) = 20.297, *p* < .001, η*G*^*2*^ = .306), but not the interaction (*F*(2, 32) = 1.688, *p* = .201, η*G*^*2*^ = .035). Multiple comparisons of the trial showed that all pairs were significant (*p*_s_ < .009).

#### Other indices

The CVs of peak times, *R*^*2*^s, and peak rates did not differ between groups in all sessions. (S Fig. 1 left, S Table 1). The transition of widths of both groups was similar to that of the stop times of the inset (S Table 2). Details of the analysis are shown in the supplemental text.

## Discussions

The results of Exp. 1 suggest that ChIs in the DS play a role in the acquisition of new duration memory. In the shift training, the peak of the response curve for the aCSF group shifted from 20 s to approximately 30 s (Fig. 2B middle). Consistent with this result, the peak and stop times of the aCSF group were higher than those of the baseline in the shift training (Fig. 2C, E). In addition, the stop times gradually increased during the shift training (inset in 2E). These results suggest that the aCSF group acquired new duration memory during the shift training. However, the peak of the response curve for the ChAT-SAP group was earlier than that of the aCSF group and similar to that of the baseline, approximately 20s (Fig. 2B left and middle). Consistent with this result, the peak and stop times of the Chat-SAP group were shorter than those of the aCSF group during the shift training (Fig. 2C, E). These results suggest that the Chat-SAP group did not acquire new duration memory. The contribution of the difference in motivational level can be excluded, because the peak rates, an index of motivation, were similar between groups (S Table 1).

The results of Exp. 1 did not clarify whether the impairment was permanent and whether relearning of the original duration memory was also impaired. To address these issues, we conducted Exp. 1B.

## Exp. 1B

All experimental procedures were approved by the Animal Research Committee of Doshisha University (A22085).

## Materials & Methods

### Subjects

The subjects are the same as those in Exp. 1A.

### Procedure

#### Additional learning (PI-40 s, sessions 33–41)

To confirm whether the impairment of acquiring new duration memory in the ChI-lesioned group was permanent, we conducted an additional nine sessions of PI-40 s training, starting three days after the test (Fig. 3A). All parameters used are the same as those in the shift training.

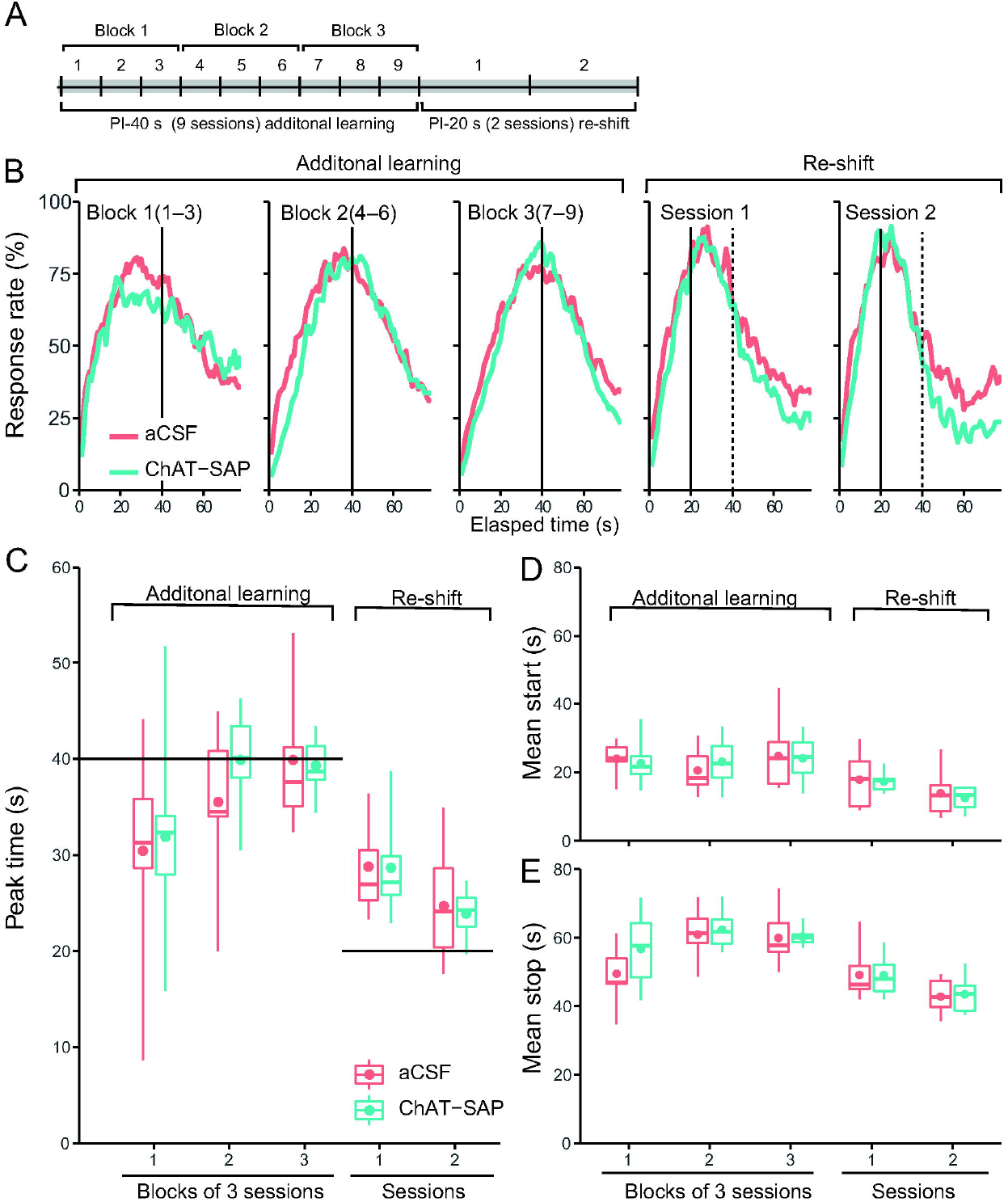

#### Re-shift training (PI-20 s, sessions 42 and 43)

To confirm whether ChI lesions impaired the utilization of already acquired duration memory, we conducted two sessions of PI-20 s training (i.e., the original required time) from the day after 41^st^ session (Fig. 3A). All parameters used are the same as those in the PI-20 s training of Exp. 1A.

## Results

Data of five rats were excluded from the analysis of Exp. 1B because the *SDs* of the fitting curves were extremely large (>80 s, the duration of an empty trial) or of poor *R*^*2*^ of fitting (below -2 *SD*) in all sessions in every block of three sessions. Thus, the numbers of subjects included were eight and nine for the ChAT-SAP and aCSF groups, respectively.

### Response distribution

Figure 3B shows mean response-rate distributions from the additional learning to the re-shift training. The peak of the aCSF group shifted to 40 s in block 1 of additional learning, whereas that of the ChAT-SAP group remained at approximately 20 s, but was flattened (Fig. 3B, 1st). From blocks 2 to 3, both distributions overlapped with peaks at approximately 40 s (Fig. 3B; 2nd, 3rd). In the re-shift training, both distributions overlapped with peaks at approximately 20 s (Fig. 3B; 4th, 5th).

### Peak time

Figure 3C shows a box and whisker plot of peak times from block 1 to re-shift training 2. In blocks 1 to 3, the peak times of both groups increased similarly (Fig. 3C). However, in the re-shift training, they decreased similarly. The mixed two-way ANOVA (group (2) × block and session (5)) showed that the main effect of the session was significant (*F*(2.35, 35.31) = 18.803, *p* < .001, η*G*^*2*^ = .446), but not the main effect of the group and the interaction (group: *F*(1, 15) = 0.193, *p* = .667, η*G*^*2*^ = .005; interaction: *F*(2.35, 35.31) = 0.543, *p* = .614, η*G*^*2*^ = .023). Multiple comparisons of sessions showed that re-shift training 1 values were lower than those in blocks 2 and 3, re-shift training 2 values were lower than those in the blocks 2 & 3 and re-shift training 1, and block 1 values were lower than those in blocks 2 and 3 (*p*_s_ < .044).

### Start time

Figure 3D shows a box and whisker plot of mean start times from block 1 to re-shift training 2. The values of both groups were similar for the additional learning (Fig. 3D). In the re-shift training, the values of both groups decreased similarly. The mixed two-way ANOVA showed that the main effect of the session was significant (*F*(4, 60) = 18.213, *p* < .001, η*G*^*2*^ = .322), but not the main effect of the group and interaction (group: *F*(1, 15) = 0.013, *p* = .910, η*G*^*2*^ = .001; interaction: *F*(4, 60) = 0.544, *p* = .704, η*G*^*2*^ = .014). Multiple comparisons of sessions showed that the values in re-shift trainings 1 and 2 were lower than those in other sessions, and that the values in re-shift training 2 was lower than that in re-shift training 1 (*p*_s_ < .038).

### Stop time

Figure 3E shows a box and whisker plot of mean stop times from block 1 to re-shift training 2. The values of both groups increased similarly in the additional learning (Fig. 3E). In the re-shift training, the values of both groups decreased similarly. The mixed two-way ANOVA showed that the main effect of the session was significant (*F*(4, 60) = 26.699, *p* < .001, η*G*^*2*^ = .538), but not the main effect of the group and interaction (group: *F*(1, 15) = 0.972, *p* = .340, η*G*^*2*^ = .022; interaction: *F*(4, 60) = 1.004, *p* = .413, η*G*^*2*^ = .042). Multiple comparisons of sessions showed that values in re-shift training 1 were lower than those in blocks 2 and 3, re-shift training 2 values were lower than those in other sessions, and block 2 values were higher than that in block 1 (*p*_s_ < .011).

### Other indices

The CVs of peak times, *R*^*2*^s, peak rates, and widths did not differ between groups (S Fig. 1 middle, S Table 3). Details of the analysis are shown in the supplemental text.

## Discussions

The results of Exp. 1B suggest that the impairment of acquisition of new duration memory by ChI lesions was transient. The peak of the response curve for the ChAT-SAP group gradually shifted toward 40s and overlapped with the curve for the aCSF group by the third block of PI-40 s training (Fig. 3B). Consistent with these results, the peak and stop times for the ChAT-SAP group also gradually increased in a way similar to that of the aCSF group (Fig. 3C, E). These findings indicate that the rats with lesions eventually acquired new duration memory.

In addition, the results of Exp. 1B suggest that relearning the original duration memory was not impaired by ChI lesions. In the re-shift training, the peak of the response curve for the ChAT-SAP group rapidly shifted toward 20 s and overlapped with the curve for the aCSF group from the first re-shift session of PI-20 s (Fig. 3B). Consistent with these results, the peak and stop times for the ChAT-SAP group also rapidly decreased in a similar way to that of the aCSF group (Fig. 3C, E). These findings indicate that lesioned rats can readily modulate their behavior depending on already acquired duration memory. Taken together, Exp. 1A and B suggest that ChI lesions have an effect on acquisition, but not utilization of once acquired duration memory or modulation of behavior depending on the change in required time.

However, the unchanged peak time of the ChAT-SAP group in Exp. 1A might be explained not only by the impairment of acquisition, but also by the co-occurrence of the normal rate of acquisition (inducing a longer peak time) and the increased speed of an internal clock (inducing a shorter peak time), assumed in the scalar expectancy theory (Gibbon, 1977). To deny the latter possibility, the effect of immunotoxic ChI lesions in the DS on the performance of PI-20 s without changing the required time between pre- and post-lesion trainings were evaluated. If the clock speed was accelerated by the lesion, then the peak time of the post-lesion session would be lower than that of the pre-lesion session.

## Exp. 2

All experimental procedures were approved by the Animal Research Committee of Doshisha University (A23078).

## Materials & Methods

### Subjects

Another cohort of fourteen male Wistar albino rats aged 3 months old were included in this experiment. They had previously undergone a guide cannula implantation surgery, PI-20 s trainings, shift training of PI-40 s, and the test. The procedures and parameters used are the same as those in Exp. 1A.

### Procedures

#### Training (PI-20 s, sessions 1–6)

The subjects underwent six sessions of PI-20 s training. (Fig. 4A). All parameters used are the same as that in the PI-20 s training in Exp. 1A. The sixth session was defined as the pre-lesion session.

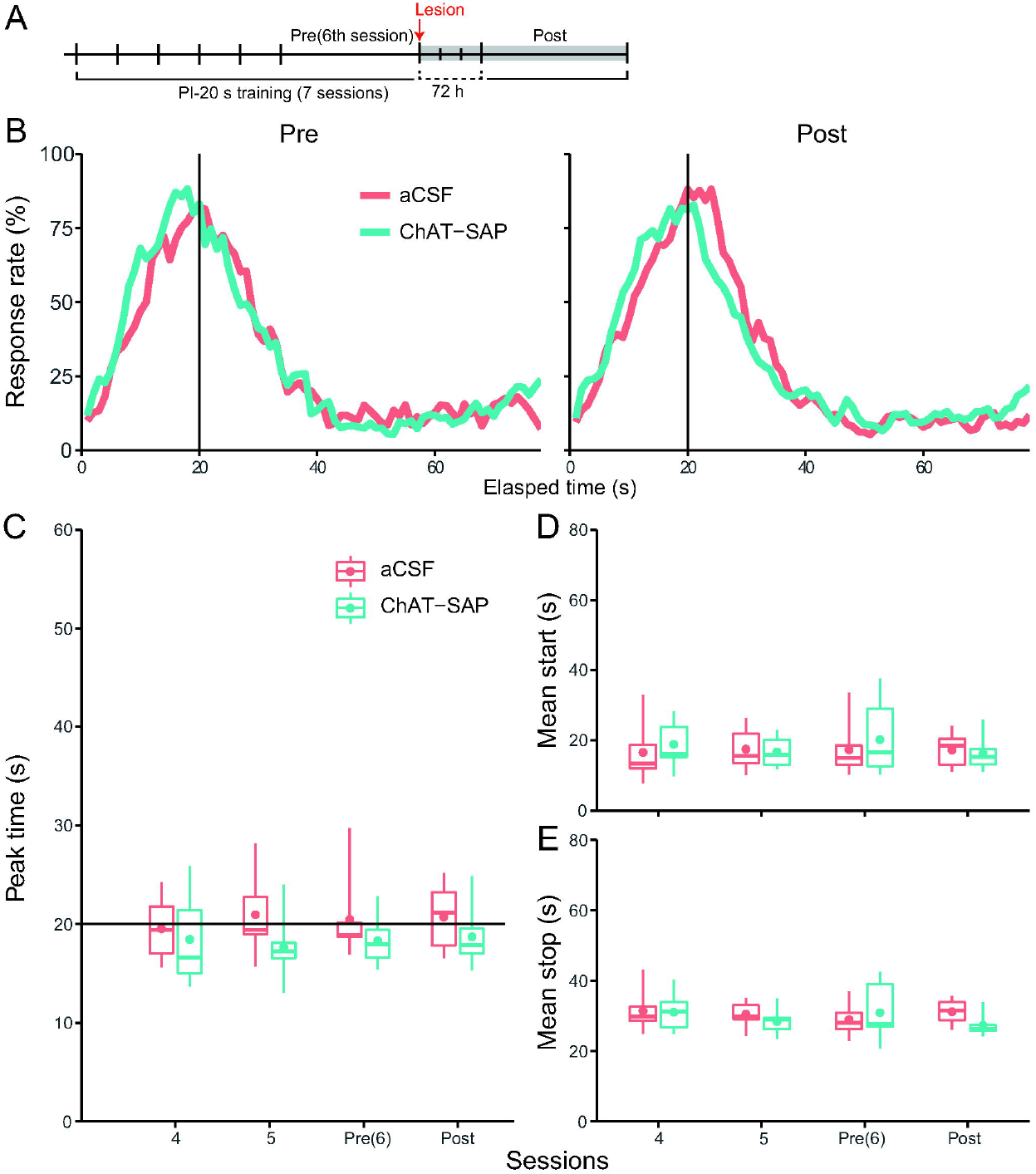

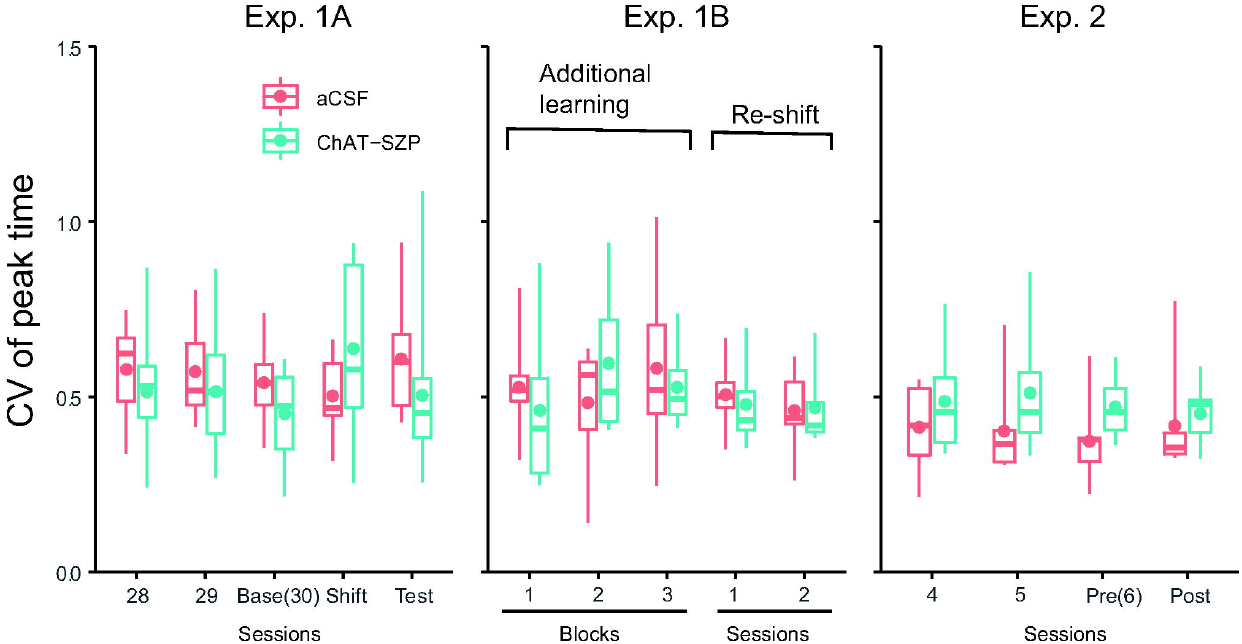

#### Immunotoxin injection

After the pre-lesion session, the subjects were injected with ChAT-SAP just as in Exp. 1A.

#### Post-lesion training (PI-20 s, session 7)

To confirm whether ChI lesions influence temporal perception, the subjects were subjected to PI-20 s training 72 h after the injection. (Fig. 4A). All parameters used are the same as those in sessions 21–30 of Exp. 1A.

## Results

### Response distribution

Figure 4B shows mean response-rate distributions in the pre- and post-lesion sessions. Both distributions overlapped with peaks at approximately 20 s (Fig. 4B).

### Peak time

Figure 4C shows a box and whisker plot of peak times from 4^th^ session to the post-lesion session. The values of both groups were at approximately 20 s in all sessions (Fig. 4C). The mixed two-way ANOVA (group (2) × session (4)) showed that none of the main effects were significant (group: *F*(1, 12) = 1.534, *p* = .239, η*G*^*2*^ = .092; session: *F*(3, 36) = 0.333, *p* = .802, η*G*^*2*^ = .006; interaction: *F*(3, 36) = 0.778, *p* = .514, η*G*^*2*^ = .014).

### Start time

Figure 4D shows a box and whisker plot of the mean start time from 4^th^ session to the post-lesion session. The values of both groups were at approximately 20 s in all sessions (Fig. 4D). The mixed two-way ANOVA showed that none of the main effects were significant (group: *F*(1, 12) = 0.074, p = .790, η*G*^*2*^ = .004; session: *F*(3, 36) = 0.483, *p* = .696, η*G*^*2*^ = .013; interaction: *F*(3, 36) = 0.657, *p* = .584, η*G*^*2*^ = .018).

### Stop time

Figure 4E shows a box and whisker plot of the mean stop time from 4^th^ session to the post-lesion session. The values of both groups were at approximately 30 s in all sessions (Fig. 4E). The mixed two-way ANOVA showed that none of the main effects were significant (group: *F*(1, 12) = 0.311, *p* = .587, η*G*^*2*^ = .014; session: *F*(3, 36) = 0.781, *p* = .513, η*G*^*2*^ = .028; interaction: *F*(3, 36) = 1.471, *p* = .239, η*G*^*2*^ = .051).

### Other indices

The CVs of peak times, *R*^*2*^s, peak rates, and widths did not differ between groups (S Fig. 1 right, S Table 4). Details of analysis are shown in the supplemental text.

## Discussions

The findings of Exp. 2 do not show that the unchanged peak time of the ChAT-SAP group in Exp. 1A might be caused by the co-occurrence of the normal rate of acquisition and the increased speed of an internal clock. If the speed of an internal clock was increased, the peak time would be shortened by the ChI lesions. However, the response curves overlapped each other in the post-lesion session. No between-group differences in the peak time, start time, and stop time in the post-lesion session were observed. Therefore, no evidence showing the speeding up of an internal clock was detected.

## General Discussions

In summary, our study suggested that ChIs play an important role in the acquisition of duration memory. In Exp. 1A, ChI lesions caused the impairment of acquisition of new duration memory. ChI lesions did not impair the utilization of once acquired duration memory or modulation of behavior depending on the change in required time in Exp. 2, interval timing itself in Exp. 3, and motivation in all experiments. Taken together, we conclude that ChI lesions delayed only the acquisition of new duration memory but did not affect the adjustment of behavior depending on changing the reinforcement schedule of PI-training and the interval timing itself.

Our findings revealed that ChI lesions in the DS are not involved in the time course of changing the peak time called “memory pattern”. In the PI training without changing the required time, the chronically systemic injection of acetylcholine receptor inhibitors gradually increased the peak time, which was maintained during the injection (Meck, 1996). This time course of change is called “memory pattern”. In a simulation by a striatal beat frequency (SBF)-ML model, a variant of the SBF model of interval timing (Matell & Meck, 2004), the time course was replicated by setting the coefficient *k\** above 1.0 (Oprisan & Buhusi, 2011). The model assumes that the DS encodes the duration and that cholinergic inputs to the DS modulate duration memory. The *k\** is believed to correspond to the synaptic level of acetylcholine in the DS (Oprisan & Buhusi, 2011). We worried that the ChI lesions in the DS might induce the “memory pattern” change in peak time because ChIs are one of the main sources of acetylcholine for the DS. However, we did not find any evidence showing that ChI lesions induced a “memory pattern”-like change in peak time (Fig. 3C). However, it is noteworthy that the cholinergic neurons they assumed were in the nucleus basalis magnocelluralis, but not ChIs. Hence, our findings would be inconsistent with that predicted by the simulation (Oprisan & Buhusi, 2011). Further studies should address the functional dissociation of acetylcholine neurons in these areas.

From the viewpoint of intra-dimensional (ID) or extra-dimensional (ED) shift of contingency (Tait et al., 2018), our suggestion was different from the roles suggested by previous studies. First, our findings suggest that ChI lesions in the DS inhibited, but did not facilitate, the ID shift task. A previous study suggested that ChI lesions in the DS enhanced reversal learning in the place discrimination task with a modified cross-maze (Okada et al., 2014). In the original learning phase of this task, a choice of one arm (location A), but not the other arm (location B), was reinforced with food. However, in the following reversal learning phase, a choice of location B, but not location A, was reinforced. Therefore, the dimension of discriminative stimulus (location of levers) was the same for both the original and reversal learning. Therefore, the task can be regarded as an ID shift task. Our task can be also regarded as an ID shift task because the dimension of the discriminative stimulus (passage of time) was unchanged, even when the required time was changed from 20 s to 40 s. However, our results suggest that ChI lesions inhibited the shift. Second, our findings provided an insight into the inhibitory role of ChI lesions, not only in the ED shift, but also in the ID shift task. In the original learning in a previous study (Aoki et al., 2015), the side of two levers (right or left) was relevant to discriminative stimulus., but the light-illuminated side located above each lever was irrelevant to one. Note that whether the right or left side of the lever and whether the light to be turned on or off were in different dimensions (i.e., extra dimension), in the following shift phase, the light-illuminated side was relevant, but the side of the levers was irrelevant. Therefore, the task can be regarded as an ED shift task. The results showed that ChI lesions in the DS inhibited this shift. However, it is difficult to directly compare our study with previous studies because many of the parameters, including the modality of discriminative stimulus (passage of time vs. allocentric/egocentric location) and the nature of variables (continuous vs. discrete) differ.

The incomplete impairment in the acquisition of new duration memory could result from the remaining ChIs. The stop time of the ChAT-SAP group gradually increased through the trials in the shift training (inset in 2E). During the additional training, the peak and stop times of the ChAT-SAP group increased similar to that of the aCSF group (Fig. 3C, E). These findings suggest that ChI lesions did not completely impair the acquisition of new duration memory. In general, an impairment in memory acquisition due to a brain lesion was permanent (Aggeleton et al., 1996; Lee & Kesner, 2004; Packard & McGaugh, 1992). Although our results contradict these studies, ChIs that respond to the presentation of reward are widespread in the striatum (Aosaki et al., 1994), and not all ChIs in the DS were lesioned (Fig. 1). Therefore, we can argue that the remaining ChIs were involved in the acquisition of new duration memory.

ChI lesions in the DS could impair the comparison between the original and new duration memory. ChIs pause their tonic firing when a reward is presented (Goldberg & Reynolds, 2011). In addition, previous studies have suggested that a pause may be involved in the encoding duration (Martel & Apicella, 2020), and occurs strongly when the predicted reward-presentation timing deviates from the actual timing (Ravel et al., 2001). ChIs may behave like the Kullback-Leibler divergence function. Therefore, we speculate that ChIs of the aCSF group paused during the shift training, and this pause encouraged the acquisition of new duration memory. Conversely, ChIs that were supposed to provide the divergence signal had been eliminated, resulting in the delayed acquisition of new duration memory in the ChAT-SAP group.

During the shift training of the aCSF group, “temporal averaging” seemed to occur. The peak time of the aCSF group was similar to the intermediate time between the required time of the baseline and shift training (i.e., 30 s). This phenomenon was also observed in previous studies (Meck et al., 1984; Hata et al., 2020; Nishioka et al., 2022). Temporal averaging is described as follows. At first, rats learned the two associations between stimuli and required times (e.g., the buzzer signals a PI of 10 s, and light signals a PI of 30 s). After that, the simultaneous presentation of these stimuli induced a peak time similar to the intermediate time between the two required times (Swanton et al., 2009). It is reasonable to assume that conflict of information between the two required times would induce the phenomena. In our study, the original and new information of duration are conflicting. Therefore, we propose that the intermediate peak time in the shift training was due to a temporal averaging.

On the premise of temporal averaging, the rats may acquire the new duration memory by “temporal scaling.” Physiological studies suggested that information on the specific duration, such as a required time, was encoded with the transition of the neural-activity pattern (Bakuhurin et al., 2017; Matell et al., 2003; Mello et al., 2015; Shikano et al., 2021; Zhou et al., 2020). A previous study argued that “temporal scaling” explains how the transition of the neural-activity pattern induces temporal averaging (Corte et al., 2022). They explain that the original neural-activity pattern encoding the original duration can also encode a longer or shorter duration than the original one by increasing or reducing the transition speed of the original neural-activity pattern (Wang et al., 2018). In other words, a neural-activity pattern can continuously, not discretely, represent durations of various lengths by changing the transition speed. Therefore, the intermediate peak time can be obtained from the intermediate transition speed (Corte et al., 2022). The above argument was supported by a recent study indicating that the subjective length of duration causally depends on the transition speed of neural-activity pattern in the DS (i.e., slowing the transition of the neural-activity pattern made the rat perceive the duration to be shorter than it was, and vice versa) (Monterio et al., 2023). In addition, temporal scaling explains how the acquisition of new duration memory was almost completed within one session, even though the original learning took dozens of sessions. The transition speed of the original neural-activity pattern may require only adjustments, rather than composing a new neural-activity pattern for the new duration memory.

Therefore, the role of ChIs might be a trigger to change the transition speed of the neural-activity pattern by providing a signal of divergence between the original duration memory and the information on the new duration. Note that they seem not to have a role in the adjustment of the transition speed of neural activity-pattern, because the re-shift of peak time to the original required time (20 s) did not (Fig. 3B, C, D, E) retard. If their role was to adjust the transition speed of it,the re-shift would be also delayed.

The role of ChIs is suggested to be different from that of M1Rs. The previous study suggested that M1R activity in the DS plays a role in the retention of the adjusted transition speed of the neural-activity pattern (Nishioka et al., 2022). Based on previous and current studies, we speculated the following serial processes from ChIs to M1Rs in the DS on the formation of duration memory. First, ChIs compare the new duration with the original duration. If ChIs detect sufficient divergence, a pause occurs. The pause then triggers unknown processes leading to an adjustment of the transition speed of the neural-activity pattern. This adjustment is retained with M1R activity.

In conclusion, our findings suggests that ChIs in the DS play an important role in the acquisition of new duration memory.

## Supporting information

STable1

STable2

STable3

STable4

## Author contributions

Masahiko Nishioka: conceptualization, software, formal analysis, investigation, writing of the original draft, writing of the review, and editing. investigation. Toshimichi Hata: methodology, software, writing the review and editing, supervision.

## Acknowledgement

We thank Fumino Fujiyama and Fuyuki Karube for teaching the immunohistological method. We thank Editage (www.editage.com) for English language editing. This work was supported by JSPS KAKENHI Grant Number JP22KJ3003 and JP22K03201.

## Data availability statement

The dataset, images of sections, and log of analysis are available from the Open Science Framework (OSF) (https://osf.io/bpfe7/?view_only=45d4707a1fce47bbaadfacbd20c980e2).

