## Supplementary material for "Cholinergic interneurons in the dorsal striatum play an important role in the acquisition of duration memory": Caption

Figure 1	**A:** The area of sections shown below (top, dotted box). Reprinted from G. Paxinos, C. Watson, The rat brain in stereotaxic coordinates. Fourth edition [CD-ROM], Copyright (1998). Photos of the representative immunohistochemically stained sections for the ChAT-SAP (middle) and aCSF (bottom) groups. Scale bar = 500 μm. Each photo was mosaiced with × 4 magnification photos. Black arrows show the closest ChIs from the cannula track. **B:** The maximum (gray) and minimum (black) areas of lesions.

Figure 2	**A:** The schedule of Exp. 1A. The gray bar shows the period after lesion occurrence. **B:** Mean response rate distributions in the baseline (left), shift (middle), and test session (right). The red line shows the aCSF group, and the blue line shows the ChAT-SAP group. Vertical solid lines show the required times for the baseline (20 s) and shift training (40 s). Dotted lines show the previous required times (20 and 40 s). **C:** Box and whisker plots of the peak times from session 28 to the test. Horizontal lines show the required times for the baseline and shift training. **D, E:** Box and whisker plots of the mean start times (D) and stop times (E) from session 28 to the test. Insets are the mean median (±*SEM*) of the start and stop times in every five trials in the shift training. Asterisks (*) indicate the significant simple main effect of the group (*p*_s_ < .050).

Figure 3	**A:** The schedule of Exp. 1B. The gray bar shows the period after lesion occurrence. **B:** Mean response rate distributions in blocks 1 to 3 of the additional training (1^st^ to 3^rd^ panels) and in session 1 and 2 of the re-shift training (4^th^ and 5^th^ panels). Red lines show the aCSF group, and blue lines show the ChAT-SAP group. Vertical solid lines show the required times for the additional training (40 s) and re-shift training (20 s). Dotted vertical lines in the re-shift session show the previous required time. **C:** Box and whisker plots of peak times from the additional training to the re-shift training. Horizontal lines show the required times for the additional training and re-shift training (40 s and 20 s, respectively). **D, E:** Box and whisker plots of the mean start times (D) and stop times from the additional training to the re-shift training.

Figure 4	**A:** The schedule of Exp. 2. **B:** Mean response rate distributions in the pre- (left) and post-lesion session (right). Red lines show the aCSF group, and blue lines show the ChAT-SAP group. Vertical solid lines show the required time (20 s). **C:** Box and whisker plots of the peak times from session 4 to the post-lesion session. Horizontal lines show the required time (20 s). **D, E:** Box and whisker plots of the mean start times (D) and stop times (E) from session 4 to the post-lesion session.

S Figure 1: Box and whisker plots of the CVs of peak times in Exp. 1A (left), 1B (middle), and Exp. 2 (right).
