## Supplemental Text for "Cholinergic interneurons in the dorsal striatum play an important role in the acquisition of duration memory"

**Exp. 1A**

***CV of peak time***: The coefficients of variations are shown in S Fig. 1 (left). The mixed two-way ANOVA showed that the interaction was significant (*F*(2.65, 42.46) = 3.338, *p* = .033, *ηG²* = .063), but not the main effects of the group and session (group: *F*(1, 16) = 0.270, *p* = .610, *ηG²* = .011; session: *F*(2.65, 42.46) = 1.104, *p* = .354, *ηG²* = .022). The simple main effect of the session in the ChAT-SAP group was significant (*F*(2.53, 22.78) = 4.291, *p* = .020, *ηG²* = .095). However, multiple comparisons of sessions in the ChAT-Sap group showed that none of the pairs were significant.

***R^2^:*** *R^2^*s of the fitting curves are shown in the S Table 1. The mixed two-way ANOVA showed that the main effect of the session was significant (*F*(2.16, 34.51) = 23.235, *p* < .001, *ηG²* = .471), but not the main effect of the group and interaction (group: *F*(1, 16) = 0.034, *p* = .857, *ηG²* = .001; interaction: *F*(2.16, 34.51) = 0.142, *p* = .882, *ηG²* = .005). Multiple comparisons of sessions showed that the values in the shift training and test were lower than those in the other sessions, and that session 28 values were lower than that at the baseline (*p*_s_ < .040).

***Peak rate:*** Peak rates are shown in the S Table 1. The mixed two-way ANOVA showed that the main effect of the session was significant (*F*(2.45, 39.28) = 24.951, *p* < .001, *ηG²* = .297), but not the main effect of the group and interaction (group: *F*(1, 16) = 0.112, *p* = .743, *ηG²* = .005; interaction: *F*(2.45, 39.28) = 0.936, *p* = .417, *ηG²* = .016). Multiple comparisons of sessions showed that the values in session 29 were higher than those in session 28, baseline, and the test, and that the shift training values were higher than those in session 28, baseline, and the test (*p*_s_ < .022).

***Width:*** Widths are shown in the S Table 1. The mixed two-way ANOVA showed that the main effect of the session was significant (*F*(2.06, 33.03) = 10.139, *p* < .001, *ηG²* = .223), but not the main effect of the group and interaction (group: *F*(1, 16) = 3.583, *p* = .077, *ηG²* = .109; interaction: *F*(2.06, 33.03) = 2.777, *p* = .075, *ηG²* = .073). Multiple comparisons of sessions showed that values in the test were lower than those in session 28 and the shift training (*p*_s_ < .023).

***Width in the shift training:*** Widths in the shift training are shown in S Table 2. The mixed two-way ANOVA showed that the main effects of the group and trial were significant (group: *F*(1, 16) = 6.560, *p* = .021, *ηG²* = .222; trial: *F*(2, 32) = 16.896, *p* = .001, *ηG²* = .242), but not the interaction (*F*(2, 32) = 2.917, *p* = .069, *ηG²* = .052). Multiple comparisons of the trial showed that all pairs were significant (*p*_s_ < .010).

**Exp. 1B**

***CV of peak time***: The coefficients of variations are shown in S Fig. 1 (middle). The mixed two-way ANOVA showed that none of the main effects were significant (group: *F*(1, 15) = 0.019, *p* = .892, *ηG²* < .001; session: *F*(2.97, 44.52) = 1.084, *p* = .365, *ηG²* = .047; interaction: *F*(2.97, 44.52) = 1.024, *p* = .391, *ηG²* = .044).

***R^2^:*** *R^2^*s of the fitting curves are shown in S Table 3. *R^2^s* of the fitting curves, peak rate, and width are shown in S Table 1. The mixed two-way ANOVA showed that the main effect of the session was significant (*F*(2.13, 32.00) = 6.412, *p* = .004, *ηG²* = .251), but not the main effect of the group and interaction (group: *F*(1, 15) = 2.472, *p* = .137, *ηG²* = .035; interaction: *F*(2.13, 32.00) = 2.009, *p* = .148, *ηG²* = .095). Multiple comparisons of sessions showed that values in block 1 were lower than that in block 3 (*p* = .014).

***Peak rate***: Peak rates are shown in S Table 3. The mixed two-way ANOVA showed that the main effect of the session was significant (*F*(4, 60) = 8.770, *p* < .001, *ηG²* = .209;), but not the main effect of the group and interaction (group: *F*(1, 15) = 3.990, *p* = .064, *ηG²* = .126; interaction: *F*(4, 60) = 0.286, *p* = .886, *ηG²* = .009). Multiple comparisons of sessions showed that values in block 1 were lower than those in the other sessions (*p*_s_ < .032).

***Width:*** Widths are shown in S Table 3. The mixed two-way ANOVA showed that the main effect of the session was significant (*F*(4, 60) = 5.459, *p* = .001, *ηG²* = .139), but not the main effect of the group and interaction (group: *F*(1, 15) = 0.350, *p* = .563, *ηG²* = .013; interaction: *F*(4, 60) = 0.962, *p* = .435, *ηG²* = .028). Multiple comparisons of sessions showed that values in re-shift training 2 were lower than those in blocks 2 and 3, and that block 2 values were higher than that in block 1 (*p*_s_ < .049).

**Exp. 2**

***CV of peak time:*** The coefficients of variations are shown in S Fig. 1 (right). The mixed two-way ANOVA showed that none of the main effects were significant (group: *F*(1, 12) = 1.627, *p* = .226, *ηG²* = .089; session: *F*(1.86, 22.33) = 0.503, *p* = .599, *ηG²* = .012; interaction: *F*(1.86, 22.33) = 0.546, *p* = .575, *ηG²* = .013).

***R^2^:*** *R^2^*s of the fitting curves are shown in S Table 4. The mixed two-way ANOVA showed that none of the main effects were significant (group: *F*(1, 12) = 0.573, *p* = .464, *ηG²* = .023; session: *F*(1.82, 21.79) = 0.776, *p* = .461, *ηG²* = .031; interaction: *F*(1.82, 21.79) = 0.047, *p* = .943, *ηG²* = .002).

***Peak rate:*** Peak rates are shown in S Table 4. The mixed two-way ANOVA showed that none of the main effects were significant (group: *F*(1, 12) = 0.585, *p* = .459, *ηG²* = .037; session: *F*(3, 36) = 1.892, *p* = .148, *ηG²* = .034; interaction: *F*(3, 36) = 0.020, *p* = .996, *ηG²* < .001).

***Width:*** Widths are shown in S Table 4. The mixed two-way ANOVA showed that none of the main effects were significant (group: *F*(1, 12) = 0.998, *p* = .338, *ηG²* = .057; session: *F*(3, 36) = 1.990, *p* < .133, *ηG²* = .043; interaction: *F*(3, 36) = 0.527, *p* = .667, *ηG²* = .012).
